# Dietary restriction promotes neuronal resilience via ADIOL

**DOI:** 10.1101/2025.06.25.661551

**Authors:** Ana Guijarro-Hernández, Shinja Yoo, George A. Lemieux, Sena Komatsu, Abdullah Q. Latiff, Rishika R. Patil, Kaveh Ashrafi

## Abstract

The steroid hormone 5-androstene-3β,17β-diol (ADIOL) was discovered in humans nearly a century ago, yet its physiological roles remain poorly defined. Here, we show that fasting and caloric restriction, two forms of dietary restriction, induce transcriptional upregulation of genes encoding CYP11A1, CYP17A1, and 17β-hydroxysteroid dehydrogenase family enzymes, promoting ADIOL biosynthesis. ADIOL, in turn, acts on the nervous system to reduce levels of kynurenic acid, a neuroactive metabolite linked to cognitive decline and neurodegeneration. This effect requires NHR-91, the *C. elegans* homolog of estrogen receptor β, specifically in the RIM neuron, a key site of kynurenic acid production. Consistent with the known benefits of fasting and caloric restriction on healthspan, enhancing ADIOL signaling improves multiple healthspan indicators during aging. Conversely, animals deficient in ADIOL signaling exhibit reduced healthspan under normal conditions and in genetic models of caloric restriction, underscoring the functional significance of this pathway. Notably, ADIOL does not significantly impact lifespan, indicating that its healthspan benefits are not simply a byproduct of lifespan extension. Together, these findings establish a physiological role for ADIOL in mediating the neuroprotective and pro-healthspan effects of fasting and caloric restriction and suggest that boosting ADIOL signaling may help narrow the gap between lifespan and healthspan. This positions ADIOL as a promising mimetic of dietary restriction effects on healthspan that could be used as a therapeutic strategy for age-related neurodegenerative conditions.

**GRAPHICAL ABSTRACT:** 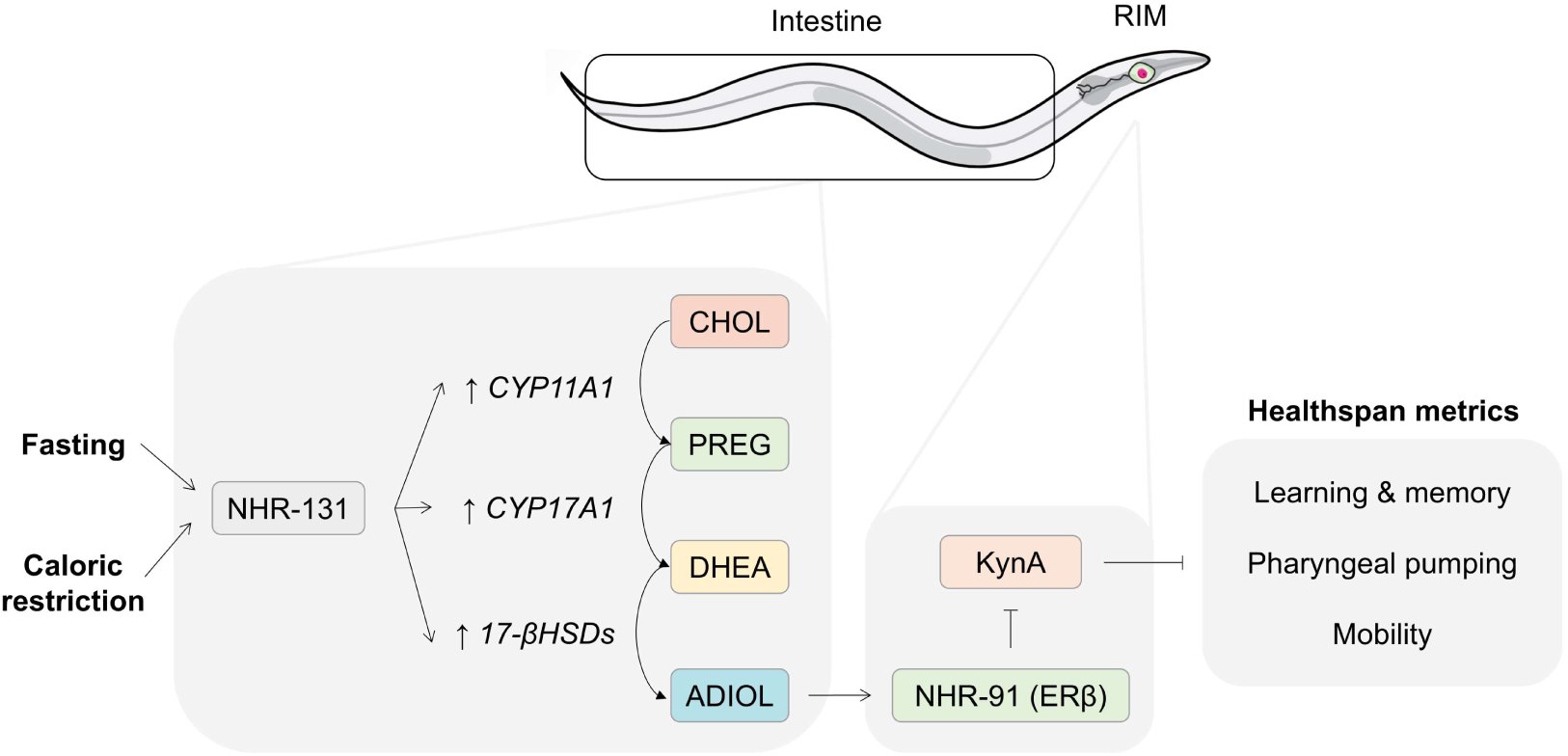

## INTRODUCTION

A central goal in aging research is to close the lifespan–healthspan gap, which refers to the difference between how long an organism lives (lifespan) and the duration over which it maintains robust physiological functions, collectively referred to as healthspan (1).

Steroid hormones are lipophilic molecules derived from cholesterol that regulate a wide range of physiological processes. Classic examples include cortisol, a glucocorticoid that modulates stress responses, metabolism, and immune function; and the sex steroids testosterone, an androgen, and 17β-estradiol (E2), an estrogen, which govern sexual development and many other biological functions (2).

Among steroid hormones, 5-androstene-3β,17β-diol (ADIOL) was discovered in humans nearly a century ago but has received limited attention and often regarded as a minor byproduct in the biosynthesis of sex steroids. ADIOL is produced from dehydroepiandrosterone (DHEA), a common precursor for testosterone and estradiol **(3; Supplementary Fig. 1a)**. In mammals, DHEA is synthesized in both the gonads and adrenal glands, and enzymatic conversions involving CYP11A1, CYP17A1, and 17β-HSDs direct it toward specific downstream products (3).

Initially classified as an androgen, ADIOL was found to possess only ∼0.21% of the androgenic activity of testosterone (4). A 1997 study showed that ADIOL binds to both estrogen receptor β (ERβ) and estrogen receptor α (ERα), with ∼3-fold higher affinity for ERβ (5), establishing it as an estrogenic compound. Unlike E2, which shows strong sex dimorphism in circulating levels, ADIOL is present at relatively similar low-nanomolar levels in males and females (6,7). Though its binding affinity is weaker than that of E2 (∼6% for ERα and ∼17% for ERβ) (5), its relatively high abundance supports its function as an endogenous ERβ ligand. Furthermore, astrocytes and microglia can locally convert circulating DHEA into ADIOL (8), suggesting a mechanism for tissue-specific regulation of ADIOL versus E2 signaling.

Crucially, ADIOL and E2 differ in their downstream effects upon binding ERβ. ADIOL, but not E2, suppresses microglial inflammation by recruiting CtBP corepressor complexes to AP-1-dependent promoters via ERβ (8), demonstrating functionally distinct transcriptional outcomes.

We recently discovered that *C. elegans* also synthesizes ADIOL (9). As in mammals, ADIOL biosynthesis in *C. elegans* depends on cytochrome P450 enzymes and 17β-HSDs (9). These enzymes are transcriptionally regulated by NHR-131, which is prominently expressed in the intestine, a metabolically active tissue in *C. elegans* (9). Elevated ADIOL levels enhance pharyngeal pumping rate, the mechanism of food intake, and improve associative learning (9). The learning paradigm employed requires *C. elegans* orthologs of NMDA and AMPA glutamate receptors, as well as CaMKII kinase, and CREB, a transcription factor, highlighting the evolutionary conservation of pathways implicated in learning and memory (10–15).

Supporting the hypothesis that ADIOL acts via ERβ-like signaling, its effects on pumping rate and learning require NHR-91, the *C. elegans* homolog of ERβ. Notably, these effects are not mimicked by E2 or testosterone (9). In *nhr-91* mutants, ADIOL has no effects on pumping or learning behaviors, but restoring wild-type *nhr-91* expression specifically in RIM neurons fully rescues ADIOL responsiveness (9). RIM neurons have previously been identified as a key site for the production of kynurenic acid (KynA), a tryptophan-derived metabolite that inhibits NMDAR-dependent activity of distinct neurons implicated in regulations of pharyngeal pumping (AVA neurons) and learning (RIM neurons) **(Supplementary Fig. 1b–c)**. We showed that ADIOL reduces KynA levels in *C. elegans*, thereby enhancing NMDAR-dependent neural functions (9). KynA similarly affects mammalian cognition: lower KynA levels are associated with improved cognition, whereas elevated KynA impairs learning and memory (16–23).

Once considered mere metabolic byproducts, kynurenine pathway (KP) metabolites such as KynA are now recognized as active signaling molecules involved in numerous pathological processes (24–29). Altered KP signaling has been implicated in neurodegenerative disorders, and modulating KP metabolites is an emerging therapeutic strategy (24,30). Additionally, KP has been linked to healthspan regulation in mammals (31). For instance, elevated kynurenine (Kyn), the precursor to KynA, is associated with sarcopenia (32), hip fractures (33,34), and cardiovascular disease (35) in older adults, and with muscle atrophy (36) and bone loss (37) in animal models.

In *C. elegans*, pharyngeal pumping and learning capacity are established behavioral readouts of healthspan (15,38). While our earlier ADIOL studies focused on day 1 adults, we have now systematically examined the effects of ADIOL during aging. Moreover, we find that ADIOL biosynthesis is induced under fasting and caloric restriction (CR)-mimicking conditions, collectively referred to as dietary restriction (DR), and that increasing ADIOL levels broadly promotes healthspan. However, unlike CR, which extends lifespan, modulating ADIOL levels causes minimal changes in lifespan. These data suggest that ADIOL mediates the neural healthspan-promoting effects of DR, independent of lifespan extension.

In summary, our findings establish ADIOL as an endogenous steroid that regulates the kynurenine pathway and neural function via ERβ/NHR-91 signaling. By enhancing healthspan without extending lifespan, ADIOL emerges as a candidate for closing the lifespan–healthspan gap.

## RESULTS

### ADIOL links nutrient status to pharyngeal pumping and learning capacity via KynA reduction

The discovery of ADIOL in *C. elegans* was facilitated by a synthetic compound, F17, initially identified through an unrelated phenotypic screen (9,39). Epistasis analyses linked F17’s activity to the kynurenine pathway metabolite kynurenic acid (KynA), and subsequent work revealed that NHR-131 and ADIOL mediate F17’s effects **(Supplementary Fig. 1b)**. Given the synthetic nature of F17 and the limited understanding of NHR-131 function, we sought to identify physiological conditions that regulate ADIOL levels.

We hypothesized that nutritional status modulates ADIOL, based on prior observations that KynA levels are nutritionally sensitive (10,40). Specifically, neural KynA levels provide a mechanism through which *C. elegans* senses nutrient availability and adjusts pharyngeal pumping accordingly (40): *C. elegans* rapidly reduce pumping upon food removal and increase it when reintroduced to food. Notably, if *C. elegans* undergo a brief 2-hour fast before refeeding, they exhibit a transient hyperactivation of pumping. Therefore, *C. elegans* increase food ingestion upon experiencing a period of fasting. This behavioral pattern is regulated by KynA levels. Fasting-induced KynA reduction leads to NMDAR-dependent activation of AVA neurons, triggering a neuropeptide Y-like signaling axis (FLP-18–NPR-5), which promotes pharyngeal pumping via serotonergic signaling (40). KynA-deficient animals show elevated pumping even when well-fed, while animals that cannot sufficiently reduce KynA levels fail to exhibit the post-fast pumping spike (40).

To determine if ADIOL contributes to fasting-induced KynA reduction, we examined pharyngeal pumping after fasting in animals lacking either NHR-91 (the ADIOL receptor) or NHR-131 (required for ADIOL biosynthesis). Unlike wild-type (WT) animals, *nhr-91* and *nhr-131* mutants failed to exhibit the hyperactivated pumping response after fasting **(Fig. 1a)**. To confirm that the phenotype was due to ADIOL deficiency, we administered exogenous ADIOL to *nhr-131* and *nhr-91* mutant animals. This fully restored the post-fasting pumping response to wild-type levels in *nhr-131* but not in *nhr-91* mutants **(Fig. 1b)**, consistent with the roles of NHR-131 and NHR-91 in ADIOL biosynthesis and response, respectively.

**Fig. 1.**
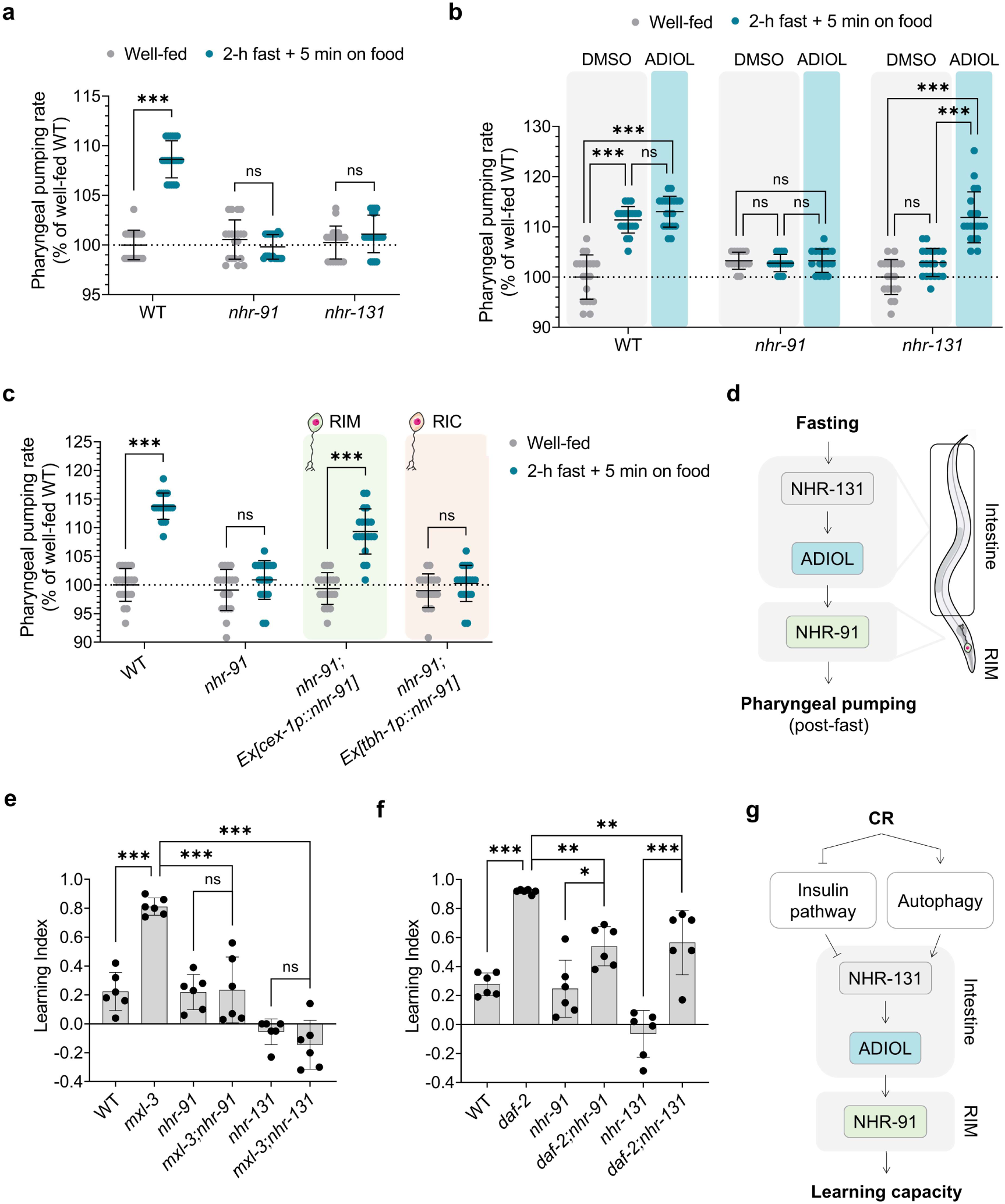
ADIOL is required for the effects of dietary restriction on pumping rate and learning capacity. **a-c**, Pharyngeal pumping rate in L4-stage animals during fed and post-fasting conditions, normalized to well-fed WT. Mean ± SD shown (n = 20 animals/condition). (a) WT, *nhr-91*, and *nhr-131* mutants. Statistics: t-test (***p < 0.001, ns: non-significant). (b) Same genotypes treated with DMSO or 10 nM ADIOL from the L1 stage. Statistics: one-way ANOVA with Bonferroni’s correction (***p < 0.001, ns: non-significant). (c) WT, *nhr-91*, and rescue strains (*nhr-91*;Ex[*cex-1p::nhr-91*,*unc-122::GFP*], and *nhr-91*;Ex[*tbh-1p::nhr-91cDNA::sl2::GFP*]) treated with DMSO or 10 nM ADIOL from the L4 stage. Statistics: t-test (***p < 0.001, ns: non-significant). **d**, Mechanism linking fasting to increased pharyngeal pumping upon re-exposure to food via ADIOL. Fasting activates NHR-131 in the intestine to promote ADIOL production, which in turn activates NHR-91 in RIM neurons, lowering KynA levels and enhancing pumping. **e**,**f**, Learning index of the caloric restriction (CR) genetic models *mxl-3* (e) and *daf-2* (f) deficient in NHR-91 or NHR-131 on day 1 of adulthood (n = 6; 15-332 animals/replicate). Mean ± SD shown. Statistics: one-way ANOVA with Bonferroni’s correction (***p < 0.001, **p < 0.01, *p < 0.05, ns: non-significant). **g**, Model illustrating the regulatory connections between CR and learning capacity via ADIOL.

NHR-91, the *C. elegans* ortholog of ERβ, is expressed in multiple neuronal types, including the RIM and RIC neurons (9), as well as some non-neural cells (41). Both RIM and RIC neurons have been shown to regulate pharyngeal pumping (42) but only RIM neurons can generate KynA (10,40). To determine the site of ADIOL action in regulation of pharyngeal pumping, we used neuron-specific promoters to express *nhr-91* exclusively in RIM or RIC neurons in *nhr-91* mutants. ADIOL increased pumping when *nhr-91* was expressed in RIM neurons, but not when expressed in RIC neurons **(Extended Fig. 1a),** pinpointing RIM neurons as the site of ADIOL action. Consistent with this, the presence of NHR-91 in RIM but not in RIC neurons was sufficient to restore the post-fasting pumping response to wild-type levels **(Fig. 1c).** Taken together, these findings support a model where fasting activates NHR-131, promoting ADIOL biosynthesis, which then binds to NHR-91 in RIM neurons to reduce KynA levels, thereby enhancing pharyngeal pumping **(Fig. 1d).**

To assess whether ADIOL’s effects are sexually dimorphic, we measured pharyngeal pumping in hermaphrodites and males treated with ADIOL or F17, which elevates ADIOL levels (9). Both compounds enhanced pumping in both sexes **(Extended Fig. 1b),** indicating that ADIOL acts independently of biological sex.

Prior work showed that CR promotes learning and memory in *C. elegans* via reductions in KynA (10). While both NMDAR-dependent and independent learning paradigms exist, the benefits of reduced KynA are specific to NMDAR-dependent learning (10). *mxl-3* mutants are a well-established CR model. Starvation represses *mxl-3* expression, which leads to increased autophagy (43). These mutants exhibit reduced KynA levels, as determined by biochemical assays, even when well-fed (10). They also show elevated pharyngeal pumping, dependent on the same signaling components required for the increased pumping observed in KynA-deficient *nkat-1* mutants **(Extended Fig. 1c),** as well as enhanced learning **(Fig. 1e)** (10). We found that the learning enhancement in *mxl-3* mutants is entirely dependent on the presence of *nhr-91* and *nhr-131* **(Fig. 1e)**. Consistent with prior data, *nhr-91* mutants show near-WT learning, while *nhr-131* mutants have significant deficits (9). These results suggest that ADIOL engagement of *nhr-91* is not required for baseline learning but is essential for learning enhancement under CR conditions recapitulated by *mxl-3* mutants. Given that *nhr-131* regulates multiple steroidogenic enzymes, including *CYP17A1/cyp-13A4*, which is likely to affect biosynthesis of multiple steroids, the more severe learning deficits of *nhr-131* mutants likely reflect its broader regulatory role in steroidogenesis.

We also examined *daf-2* (insulin receptor-deficient) and *rict-1* (mTOR pathway-deficient) mutants because modulations of insulin and mTOR pathways also mimic aspects of CR, and the *daf-2* and *rict-1* mutants were previously shown to exhibit reduced KynA levels and enhanced learning (10). Losses of *nhr-91* and *nhr-131* partially suppressed learning enhancement in *daf-2* mutants **(Fig. 1f)**, whereas loss of *nhr-91* had no effect on the learning phenotype of *rict-1* mutants (**Extended Fig. 1d**). These findings support the idea that ADIOL plays a role in conditions of CR to increase learning capacity. They also reveal that reductions of KynA levels can be achieved independently of ADIOL signaling.

### ADIOL levels increase under fasting and CR conditions through the upregulation of *CYP11A1*, *CYP17A1* and *17-βHSDs* orthologs

ADIOL is synthesized from cholesterol via a well-characterized enzymatic cascade: CYP11A1 converts cholesterol to pregnenolone (PREG); CYP17A1 transforms PREG into dehydroepiandrosterone (DHEA); and 17β-hydroxysteroid dehydrogenases (17β-HSDs) convert DHEA to ADIOL **(Fig. 2a)**. Prior work showed that the synthetic compound F17 increases ADIOL levels in *C. elegans* by activating NHR-131, which upregulates multiple *CYP17A1* and *17β-HSD* homologs **(Fig. 2a)** (9). To understand how ADIOL levels rise during fasting or in CR models, we analyzed the expression of these NHR-131-dependent, F17-upregulated genes after 2 hours of fasting and in *daf-2* and *mxl-3* mutants using RT-qPCR. We also included *cyp-44A1*, the homolog of *CYP11A1*, which catalyzes the first step in the pathway, as its upregulation, though not induced by F17, could contribute to fasting/CR-induced ADIOL synthesis.

**Fig. 2.**
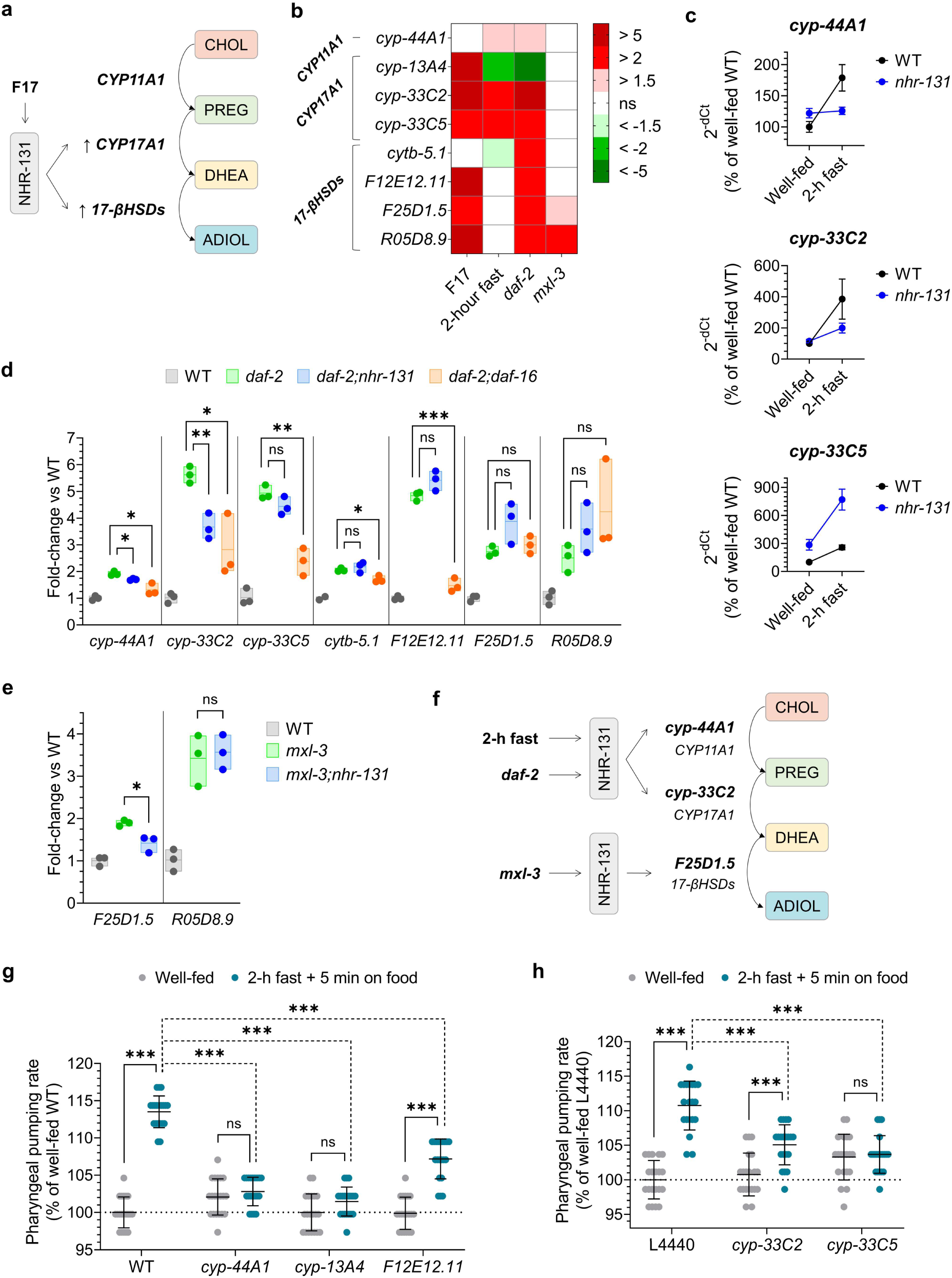
Fasting and genetic models of CR upregulate ADIOL biosynthesis genes. **a**, F17-induced activation of NHR-131 promotes ADIOL biosynthesis from cholesterol by upregulating *C. elegans* homologs of human CYP17A1 and 17-βHSDs, which catalyze the conversion of pregnenolone (PREG) to dehydroepiandrosterone (DHEA), and DHEA to ADIOL, respectively (9). **b**, RT-qPCR analysis of *CYP11A1*, *CYP17A1*, and *17β-HSD* homologs in L4 animals treated with 2.5 µM F17 or under conditions shown (n = 3; ∼300 animals/replicate). Mean ± SD values shown relative to DMSO-treated controls (F17-treated animals) or to well-fed WT animals (2-hour fasting and the mutants shown). Statistics: one-way ANOVA with Bonferroni’s post hoc test on ΔCt values. Non-statistically significant differences (ns) are shown in white. **c**, RT-qPCR of *cyp-44A1*, *cyp-33C2*, and *cyp-33C5* in WT and *nhr-131* L4 animals under fed and fasting conditions (n = 3; ∼300 animals/replicate). Mean ± SD values shown relative to well-fed WT. Statistics: two-way ANOVA on ΔCt values (positive interaction for *cyp-44A1* (p = 0.0004) and *cyp-33C2* (p = 0.02); no interaction for *cyp-33C5* (p = 0.89)). **d**,**e**, RT-qPCR analysis of a subset of ADIOL biosynthesis genes in *daf-2* (d) and *mxl-3* (e) mutants deficient in NHR-131 or DAF-16 (n = 3; ∼300 animals per condition). Mean ± SD values shown relative to WT. Statistics: t-test on ΔCt values (***p < 0.001, **p < 0.01, *p < 0.05, ns: not significant). **f**, Model showing the dependence on *nhr-131* for the upregulation of ADIOL biosynthetic genes in the context of fasting and CR. **g**,**h**, Pharyngeal pumping rate in L4-stage animals under fed and post-fasting conditions (n = 20 animals/condition). Mean ± SD values shown relative to well-fed WT. Statistics: one-way ANOVA with Bonferroni’s correction (***p < 0.001, ns: non-significant). (g) WT, *cyp-44A1*, *cyp-13A4*, and *F12E12.11* animals were analyzed. (h) WT animals exposed to RNAi clones L4440, *cyp-33C2*, or *cyp-33C5* from the L1 stage.

A subset of *CYP11A1* and *CYP17A1* homologs was upregulated after 2 hours of fasting and in *daf-2* mutants. Specifically, both fasting and loss of *daf-2* induced expression of *cyp-44A1* (*CYP11A1*), and *cyp-33C2* and *cyp-33C5* (*CYP17A1* homologs) **(Fig. 2b)**. Analysis of their expression in *nhr-131*–deficient animals revealed that the fasting-induced changes in *cyp-44A1* and *cyp-33C2* expression are dependent on *nhr-131* **(Fig. 2c)**, and in the case of *daf-2* mutants, also dependent on *daf-16*, the main downstream effector of *daf-2* **(Fig. 2d)**. All analyzed *17β-HSD* homologs showed elevated expression in *daf-2* mutants **(Fig. 2b)**, whereas in *mxl-3* mutants only *F25D1.5* and *R05D8.9* exhibited increased expression **(Fig. 2b)**. Among these, only the upregulation of *F25D1.5* in *mxl-3* mutants was *nhr-131*–dependent **(Fig. 2e)**. These findings indicate that fasting and CR upregulate genes involved in ADIOL synthesis, in some cases via *nhr-131* **(Fig. 2f)**.

We reasoned that if these biosynthetic genes mediate ADIOL production, their loss should mimic the effects of impaired ADIOL signaling. To test this, we examined post-fast pharyngeal pumping, a behavior known to require ADIOL and *nhr-91* **(Fig. 1)**. *cyp-44A1* mutants failed to elevate pharyngeal pumping following refeeding, while *cyp-13A4* and *F12E12.11* mutants showed only partial responses **(Fig. 2g)**. Similarly, RNAi knockdown of *cyp-33C2* and *cyp-33C5* (*CYP17A1* homologs) failed to fully activate pumping upon post-fast refeeding **(Fig. 2h)**. These partial requirements likely reflect enzymatic redundancy, with multiple enzymes capable of catalyzing each step.

Together, these findings support a model in which fasting and CR enhance ADIOL production through upregulation of key biosynthetic genes, *CYP11A1*, *CYP17A1* and *17-βHSDs* orthologs.

### ADIOL promotes neuronal healthspan during aging through KynA reduction

A previous study demonstrated that ADIOL enhances learning and memory in aged animals by lowering KynA levels (11). However, whether ADIOL is required to maintain normal learning capacity throughout aging remains unknown. To address this, we examined learning capacity in *nhr-91* and *nhr-131* mutants across aging. While wild-type animals exhibit age-associated declines in learning, *nhr-91* mutants, which are unable to respond to ADIOL, showed a markedly earlier and more pronounced decline **(Fig. 3a)**. Similarly, *nhr-131* mutants, which lack ADIOL biosynthesis, displayed severe impairments from early adulthood **(Fig. 3a)**. The more pronounced phenotype of *nhr-131* mutants is consistent with the notion that this transcription factor is likely to regulate generation of steroids beyond ADIOL.

**Fig. 3.**
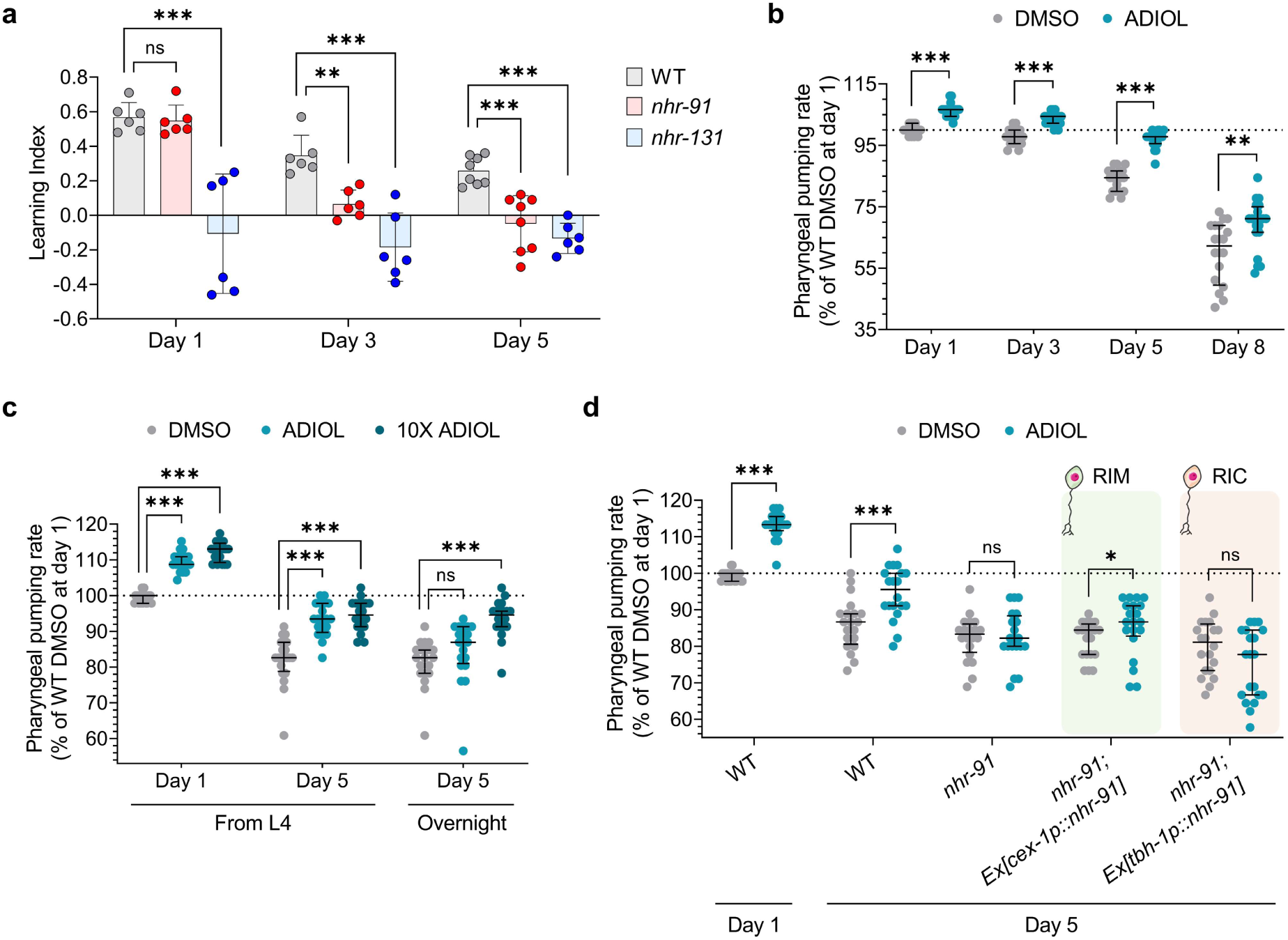
ADIOL promotes neuronal healthspan during aging through KynA reduction. **a**, Learning index of WT, *nhr-91* and *nhr-131* animals on day 1, 3, and 5 of adulthood (n = 6-8; 25-232 animals/replicate) after exposure to 50 µM FUDR from day 1. Mean ± SD shown. Statistics: one-way ANOVA with Bonferroni’s correction (***p < 0.001, **p < 0.01, ns: non-significant). **b**-**d**, Pharyngeal pumping rate during aging (n = 20 animals/condition). Animals were exposed to 50 µM FUDR from day 1. Median ± IQR values shown relative to WT DMSO-treated day 1 adults. (b) WT animals treated with DMSO or 10 nM ADIOL from the L4 stage, assayed on days 1, 3, 5, and 8 of adulthood. Statistics: t-test (for day 1 and day 3 adults) or Mann-Whitney U test (for day 5 and day 8 adults) (***p < 0.001, **p < 0.01). (c) WT animals treated with DMSO, 10 nM ADIOL or 100 nM ADIOL (10X) on days 1 and 5 of adulthood. For day 5 adults, treatment began either at the L4 stage or on day 4 (overnight). Statistics: one-way ANOVA with Bonferroni’s correction (for day 1 adults) or Mood’s median test followed by multiple comparisons (for day 5 adults) (***p < 0.001, ns: non-significant). (d) Day 5 adults of WT, *nhr-91*, *nhr-91*;Ex[*cex-1p::nhr-91,unc-122::GFP*], and *nhr-91*;Ex[*tbh-1p::nhr-91cDNA::sl2::GFP*] treated with DMSO or 10 nM ADIOL from the L4 stage. Statistics: t-test or Mann-Whitney U test depending on distribution (***p < 0.001, *p < 0.05, ns: non-significant).

Given that pharyngeal pumping is another established healthspan readout, we also assessed this behavior in *nhr-91* and *nhr-131* mutants during aging. While the pumping rate of wild type animals declined by 15% between day 1 and day 5 adults, *nhr-91* and *nhr-131* mutants exhibited 20% declines during the same period **(Extended Fig. 2a)**. To test whether ADIOL’s effects on pumping persist with age, we treated wild-type animals with ADIOL from the L4 stage onward. While pharyngeal pumping declined with age in untreated animals, ADIOL treatment blunted this decline **(Fig. 3b)**. To determine whether administration of ADIOL once the decline has begun is also effective, we treated day 4 adult animals with either the standard or a tenfold higher ADIOL dose and measured pumping on day 5. While the standard dose had no effect, the higher dose restored pumping to levels comparable to those seen with lifelong treatment **(Fig. 3c)**. Day 5 animals were previously demonstrated to have significantly elevated levels of KynA (11), providing a potential rationale as to why elevated doses of ADIOL are needed to elicit an increase in pharyngeal pumping.

To determine whether ADIOL’s effects on pumping remain dependent on RIM neuron in aged animals, we examined *nhr-91-*deficient animals in which *nhr-91 was* specifically expressed in RIM or RIC neurons. Only RIM-specific reconstitution restored the ability of ADIOL to enhance pharyngeal pumping in day 5 adults, confirming the continued importance of this neuronal locus **(Fig. 3d)**.

To examine other indicators of healthspan, we next assessed mobility by measuring thrashing, the swimming movement of worms in liquid. As expected, thrashing declined with age **(Fig. 4a)**. The age-dependent declines in thrashing rate and increases in its variability were similar to previously reported measurements of thrashing (44,45). ADIOL treatment from the L4 stage increased thrashing in day 1 adults, but its effects at later ages were modest and potentially obscured by the high variability of thrashing rate in the absence of treatment **(Fig. 4a)**. To better assess age-related effects, we repeated the experiment without FUDR and using a tenfold higher ADIOL concentration. Under these conditions, ADIOL robustly enhanced thrashing in aged animals **(Fig. 4b)**. However, in the absence of FUDR and ADIOL, we observed an unexpected increase in thrashing rate of day 5 wild-type animals, coincident with termination of the egg-laying period **(Fig. 4b)**. The basis for this increase is unclear.

**Fig. 4.**
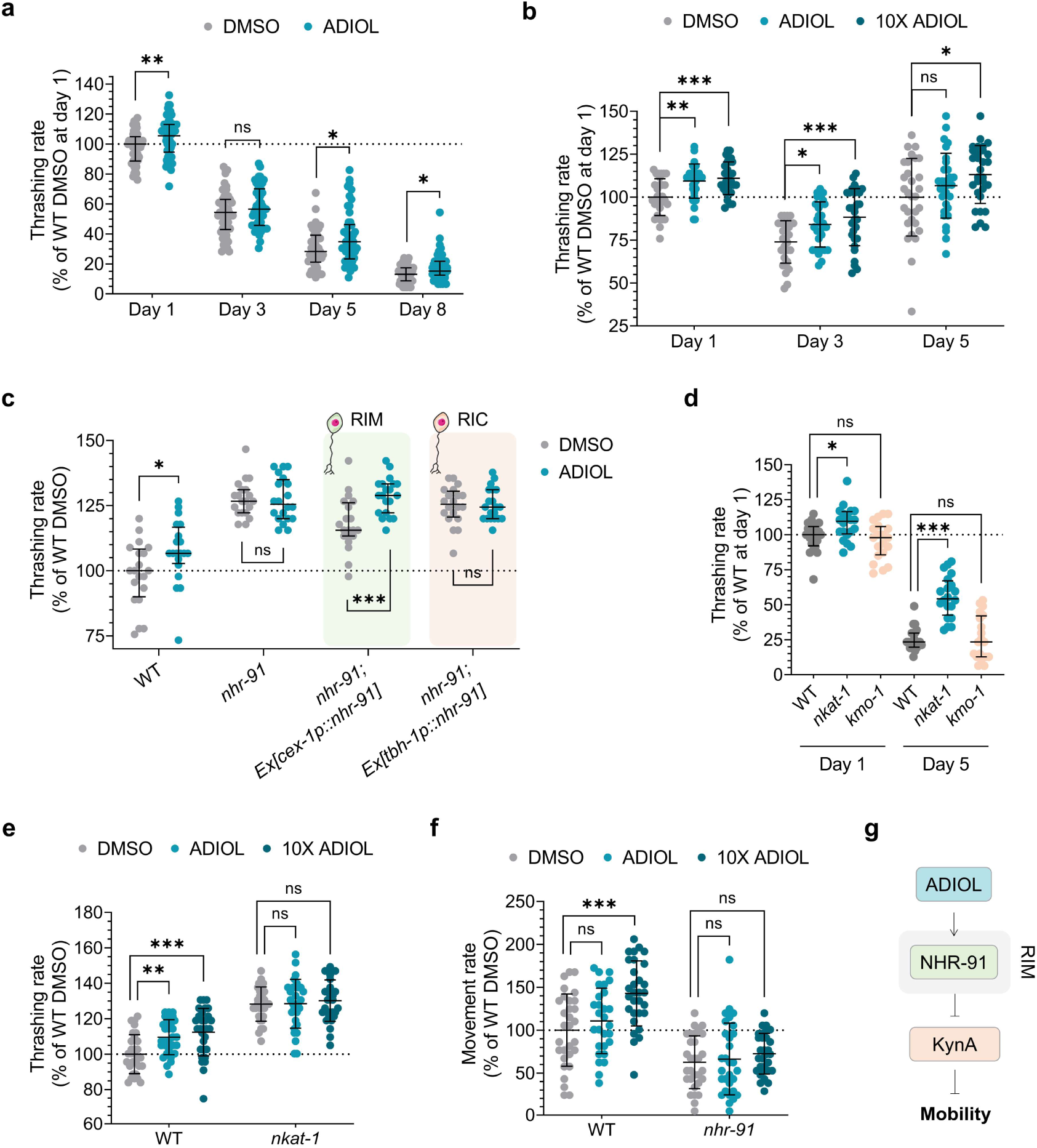
ADIOL promotes mobility during aging through KynA reduction. **a**-**d,** Thrashing rate during aging (n = 20-50 animals/condition). Unless otherwise indicated, median ± IQR values shown relative to WT DMSO-treated day 1 adults, and animals were exposed to 50 µM FUDR starting on day 1 and treated with DMSO or ADIOL from the L4 stage. (a) WT animals treated with DMSO or 10 nM ADIOL on days 1, 3, 5, and 8 of adulthood. Statistics: t-test (days 1, 3) or Mann-Whitney U test (days 5, 8) (**p < 0.01, *p < 0.05, ns: non-significant). (b) WT animals treated with DMSO, 10 nM ADIOL or 100 nM ADIOL (10X) on days 1, 3, and 5. No FUDR was used in this assay. Mean ± SD values shown relative to WT DMSO-treated day 1 adults. Statistics: one-way ANOVA with Bonferroni’s correction (***p < 0.001, **p < 0.01, *p < 0.05, ns: non-significant). (c) Day 5 adults of WT, *nhr-91*, *nhr-91*;Ex[*cex-1p::nhr-91,unc-122::GFP*], and *nhr-91*;Ex[*tbh-1p::nhr-91cDNA::sl2::GFP*] treated with DMSO or 10 nM ADIOL. Statistics: t-test (***p < 0.001, *p < 0.05, ns: non-significant). (d) WT, *nkat-1* and *kmo-1* animals on day 1 and 5 of adulthood. Statistics: one-way ANOVA with Bonferroni’s correction (***p < 0.001, *p < 0.05, ns: non-significant). **e**, Thrashing rate of WT and *nkat-1* animals on day 1 of adulthood after DMSO, 10 nM ADIOL or 100 nM ADIOL (10X) starting at L4 stage (n = 20 animals/condition). Data are shown as percentage relative to WT DMSO-treated animals (mean ± SD). Statistics: one-way ANOVA with Bonferroni’s correction (***p < 0.001, **p < 0.01, ns: non-significant). **f**, Spontaneous movement of WT and *nhr-91* animals on day 1 of adulthood after treatment with DMSO, 10 nM ADIOL or 100 nM ADIOL (10X) starting at the L4 stage (n = 20 animals/condition). Data are shown as percentage relative to WT DMSO-treated animals (mean ± SD). Statistics: one-way ANOVA with Bonferroni’s correction (***p < 0.001, ns: non-significant). **g**, Model summarizing the effect of ADIOL on mobility.

We next tested whether ADIOL’s effects on thrashing also require NHR-91 in RIM neurons. As with pharyngeal pumping, ADIOL enhanced thrashing only when NHR-91 was reconstituted in RIM neurons **(Fig. 4c)**. We next asked whether ADIOL’s effect on thrashing depends on KynA. *nkat-1* mutants (low KynA) (9,40) exhibited elevated thrashing on both day 1 and day 5, while *kmo-1* mutants (high KynA) (9,40) did not further diminish the already reduced thrashing of day 5 animals **(Fig. 4d)**. Consistent with a KynA-dependent mechanism, ADIOL failed to further enhance thrashing in *nkat-1* mutants **(Fig. 4e)**.

Analysis of spontaneous animal movement on solid media also demonstrated that ADIOL treatment increases locomotion in an NHR-91–dependent manner **(Fig. 4f)**, further validating the notion that ADIOL broadly improves healthspan **(Fig. 4g)**.

We also evaluated whether ADIOL influences other aging-related phenotypes. ADIOL-treated animals showed a trend towards modestly increased resistance to osmotic stress (NaCl) **(Extended Fig. 2b)**. A similar trend, albeit not reaching statistical significance, was seen in *nkat-1* mutants (**Extended Fig. 2c**). Moreover, chemotaxis to benzaldehyde was unaffected by ADIOL at any age **(Extended Fig. 2d)**. These data indicate that ADIOL enhances multiple, but not all, indicators of healthspan.

Together, these findings show that loss of ADIOL signaling via NHR-91 accelerates decline in multiple healthspan indicators while exogenous ADIOL administration mitigates these declines during aging through a mechanism that is mimicked by KynA depletion.

### The beneficial effects of ADIOL on healthspan are not merely a result of lifespan extension

An important question raised by these findings was whether ADIOL’s healthspan benefits simply reflect changes in lifespan. Animals lacking the ADIOL receptor (*nhr-91* mutants) showed a modest reduction in mean lifespan (approximately one day shorter than wild-type animals) while *nhr-131* mutants, which likely lack biosynthesis of multiple steroid hormones including ADIOL, exhibited a more pronounced decrease **(Fig. 5a)**. However, ADIOL treatment did not extend the lifespan of wild-type animals **(Fig. 5b)**. Furthermore, CR extended the lifespan of *nhr-91* mutants, and loss of *nhr-91* had no effect on the extended lifespans of long-lived *daf-2*, *mxl-3*, or *eat-2* mutants **(Fig. 5c–f)**.

**Figure 5.**
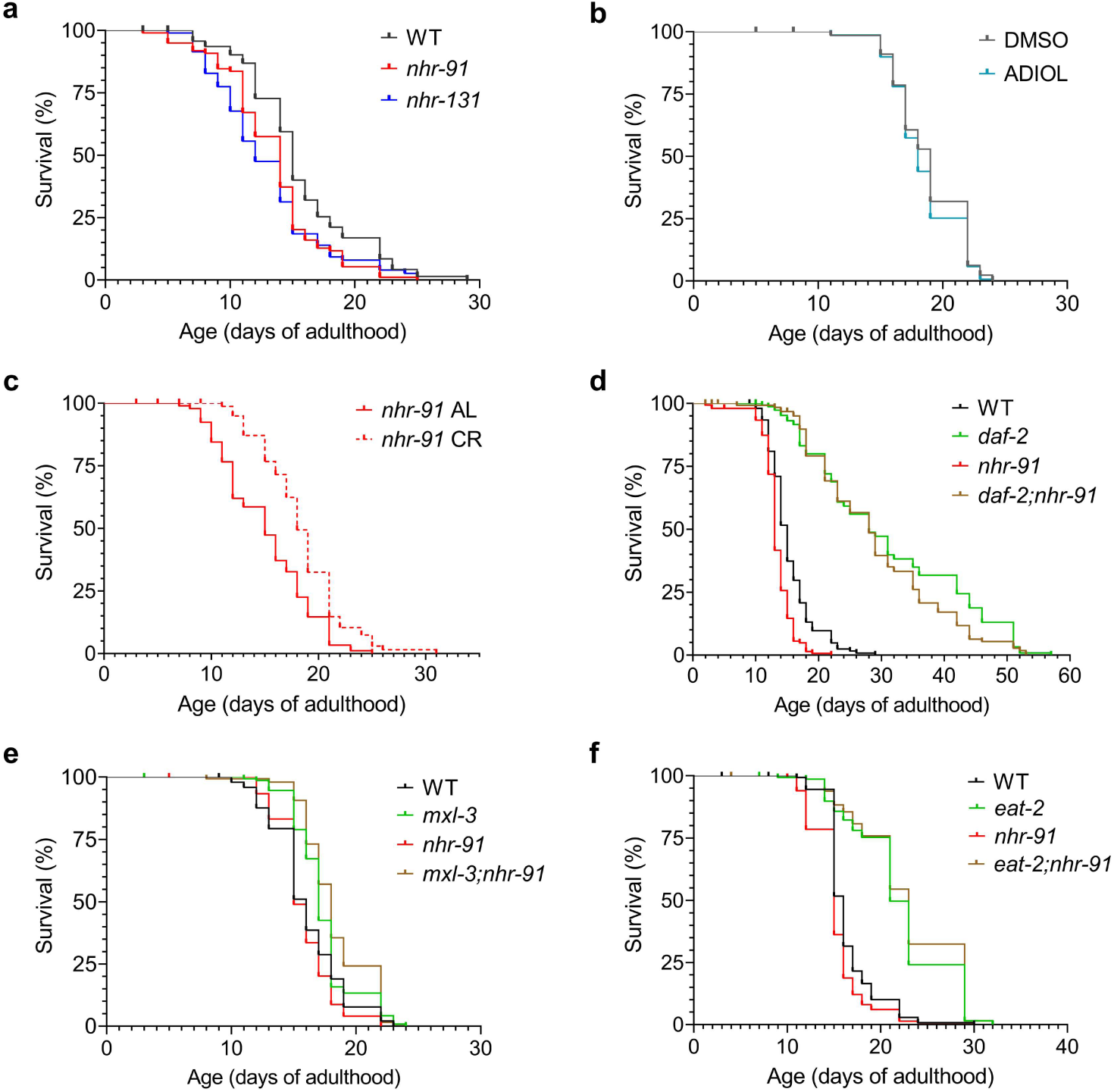
ADIOL has no impact on longevity. **a**, Lifespan of WT, *nhr-91*, and *nhr-131* animals at 20 °C. Median survival: 15 days (WT, n = 100), 14 days (*nhr-91*, n = 99), and 12 days (*nhr-131*, n = 96). Log-rank test with Bonferroni correction showed significant differences between WT and *nhr-91* (p = 0.002) and between WT and *nhr-131* (p = 0.001). **b**, Lifespan of WT animals treated with DMSO or 10 nM ADIOL from the L4 stage. Median survival: 19 days (DMSO, n = 136) and 18 days (ADIOL, n = 150). No significant differences were detected (log-rank test, p > 0.01). **c**, Lifespan of *nhr-91* animals fed *ad libitum* (AL) or under CR. Median survival: 15 days (AL, n = 100) and 18 days (CR, n = 100). CR significantly extended lifespan (log-rank test, p < 0.001). **d-f**, Lifespan curves of the CR genetic models *daf-2* (d) *mxl-3* (e) and *eat-2* (f) deficient in NHR-91. Kaplan–Meier survival curves were compared using the log-rank test with Bonferroni correction. (d) Median survival was 15 days (WT, n = 154), 28 days (*daf-2* ,n = 150), 13 days (*nhr-91*, n = 151), and 28 days (*daf-2;nhr-91*, n = 150). No significant differences between *daf-2* and *daf-2;nhr-91* mutants (p > 0.01). (e) Median survival was 16 days (WT, n = 150), 17 days (*mxl-3*, n = 150), 15 days (*nhr-91*, n = 150), and 18 days (*mxl-3;nhr-91*, n = 150). No significant differences between *mxl-3* and *mxl-3;nhr-91* mutants (p > 0.01). (f) Median survival was 16 days (WT, n = 150), 21 days (*eat-2*, n = 150), 15 days (*nhr-91*, n = 150), and 23 days (*eat-2;nhr-91*, n = 150). No significant differences between *eat-2* and *eat-2;nhr-91* mutants (p > 0.01).

These findings demonstrate that ADIOL’s healthspan-promoting effects are not secondary to changes in lifespan. This conclusion is further supported by previous studies showing that reductions in KynA levels do not alter lifespan (10).

## DISCUSSION

Our results indicate that ADIOL biosynthesis is enhanced under fasting and CR conditions, and that ADIOL signaling through the *C. elegans* homolog of estrogen receptor β, NHR-91, mediates several healthspan benefits associated with fasting and CR. ADIOL’s effects on pharyngeal pumping, learning capacity, and thrashing, all indicators of healthspan, are dependent on NHR-91 activity in RIM neurons, a primary site of KynA production. Consistent with the KynA-lowering action of ADIOL, fasting/CR reduces KynA levels (10), and animals deficient in KynA exhibit improved performance in the same healthspan indicators. While increasing ADIOL levels promotes healthspan during aging, loss of *nhr-91* accelerates age-associated decline in the specific indicators of healthspan tested. Together, these findings provide the first evidence that the steroid hormone ADIOL mediates several neural benefits of fasting and CR.

Supporting the notion that ADIOL biosynthesis is nutritionally regulated, several genes predicted to encode enzymes involved in its synthesis (homologs of *CYP11A1*, *CYP17A1*, and *17β-HSDs*) are upregulated following short-term fasting or in genetic models that mimic aspects of CR (**Fig. 2**). These enzymes convert cholesterol to ADIOL via pregnenolone (PREG) and DHEA (3).

While the link between fasting/CR and ADIOL production is a novel insight, prior studies in other species are consistent with our findings. Plasma DHEA levels increase during fasting in male zebra finches and in the lizard *Anolis sagrei* (46,47), and in rodents, fasting induces hepatic CYP17A1 expression and elevates DHEA levels via PGC-1α activation (48,49). In parallel, prolonged CR in mice increases expression of estrogen receptors, including ERβ (50), and in humans, CR reduces circulating kynurenine, the precursor of KynA (51). However, whether ADIOL levels are modulated by fasting/CR in mammals remains unknown.

We found that not all CR models depend on ADIOL to improve learning capacity **(Fig. 1e–f, Extended Fig. 1d)**, suggesting that while ADIOL is a key contributor, it is not the sole mediator of CR-induced benefits. We previously found that fasting and CR models regulate *kmo-1*, encoding kynurenine monooxygenase-1, which competes with KATs for the substrate kynurenine (10). Thus, KMO-1 activity may influence KynA production independently of ADIOL, highlighting multiple regulatory routes by which fasting/CR can lower KynA and improve neural function.

Importantly, ADIOL did not extend lifespan in wild-type animals (**Fig. 5**), and its receptor mutant, *nhr-91*, showed only a modest lifespan reduction. Furthermore, *nhr-91* was not required for lifespan extension in long-lived mutants such as *daf-2*, *mxl-3*, or *eat-2*. These findings indicate that ADIOL’s healthspan-promoting effects are not indirect consequences of lifespan extension. This distinction is especially relevant given the growing interest in closing the lifespan-healthspan gap (1).

What molecular mechanisms underlie the ADIOL–NHR-91/ERβ–KynA axis in promoting neural resilience? While the full picture is not yet known, KynA’s signaling roles offer a plausible mechanistic framework. We previously showed that the pharyngeal pumping and learning benefits of KynA reduction in *C. elegans* depend on enhanced activity of specific neurons in an NMDAR-dependent manner (10). However, NMDAR-independent mechanisms are also plausible since, in mammals, KynA is also a known agonist of the aryl hydrocarbon receptor (AhR) and GPR35, a GPCR (52–54).

Together, our findings show that ADIOL biosynthesis pathway is nutritionally sensitive and that this overlooked steroid hormone improves multiple indicators of neural healthspan and resilience. The results also point to KynA reducing effects of ADIOL as the likely molecular explanation. In light of these findings, ADIOL emerges as a promising mimetic of dietary restriction effects on healthspan, highlighting its potential as a therapeutic strategy for age-related neurodegenerative conditions.

## MATERIALS AND METHODS

### Strains and maintenance

Strains used in this study are listed in **Supplementary Table 1**. Plasmid constructs were generated by standard PCR amplification from *C. elegans* genomic DNA or cDNA and assembled using NEBuilder HiFi DNA Assembly (New England Biolabs). Transgenic extrachromosomal arrays included Ex[*cex-1p::nhr-91,unc-122::GFP*], which used a 0.9 kb region upstream of the ATG start codon of *cex-1* as promoter and included *unc-122::GFP* as a coinjection marker. Ex[*tbh-1p::nhr-91cDNA::SL2::GFP*] used 3,455 bp upstream of the ATG of *tbh-1* as promoter and contained the full *nhr-91* cDNA followed by a bicistronic SL2::GFP reporter. The 3′ untranslated region (UTR) of *unc-54* was used for all constructs to ensure proper transcript processing. Transgenic animals carrying non-integrated extrachromosomal arrays were generated by microinjecting expression plasmids into the gonad arm at final concentrations ranging from 5 to 50 ng/µL. Double mutants were obtained through standard genetic crosses between males and hermaphrodites.

Unless otherwise specified, *C. elegans* were maintained at 20 °C on 6-cm NGM agar plates seeded with *Escherichia coli* OP50, following established protocols (55).

### RNAi

RNAi clones targeting *cyp-33C2* and *cyp-33C5* (Ahringer library) (56) were used for gene knockdown experiments. The empty vector L4440 was used as a negative control, and the *nkat-1* RNAi clone was included as a positive control. *E. coli* HT115 strains containing the respective RNAi plasmids were grown overnight at 37 °C in 5 mL of LB supplemented with 100 µg/mL carbenicillin. Cultures were then diluted 1:40 into 5 mL of fresh LB with 100 µg/mL carbenicillin and grown at 37 °C with shaking until reaching an OD_600_ of 0.4-0.6. Double-stranded RNA production was induced by adding 1 mM IPTG and incubating for 4 h at 37 °C. To spike the induction, cultures were supplemented with an additional 100 µg/mL carbenicillin and 1 mM IPTG. A total of 250 µL of induced culture was seeded onto 6-cm NGM plates containing 100 µg/mL carbenicillin and 1 mM IPTG. Plates were incubated overnight at room temperature to allow bacterial lawn formation. Synchronized L1 animals were then transferred onto the RNAi plates and maintained at 20 °C until the L4 stage for further experiments.

### Pharmacological treatments

The pharmacological treatments used in this study were F17 (custom synthesis, TimTec) and 5-androstene-3β,17β-diol (ADIOL) (Steraloids). Both compounds were dissolved in DMSO, which served as the vehicle. F17 was applied at a final concentration of 2.5 μM, and ADIOL was administered at a final concentration of 10 nM, with a subset of experiments including a tenfold higher dose (10X; 100 nM) to assess dose-dependent effects. Concentrations were calculated based on the total NGM volume per plate and were mixed with *E. coli* OP50 prior to seeding. Unless otherwise specified, treatments were initiated at the L4 larval stage.

### 2-h fasting

Fasting was conducted by collecting animals in 1 mL of S-basal + 0.05% PEG-8000 and washing them three times with the same solution. After washing, worms were rotated for 2 hours in the same buffer. For pharyngeal pumping assessment, animals were transferred to plates seeded with *E. coli* OP50 and allowed to acclimate for at least 5 minutes before counting.

### Pharyngeal pumping

Pharyngeal pumping was assessed by counting the number of contractions of the posterior pharyngeal bulb over a 10-second period, as previously described (40). The assay was performed on 20 L4 or day 1 adult animals per condition. Animals were age-synchronized using standard hypochlorite treatment of gravid hermaphrodites to isolate eggs, which were allowed to hatch overnight in S-basal supplemented with 0.05% PEG-8000. Approximately 150 synchronized L1 larvae per condition were transferred onto 6-cm NGM plates and incubated at 20 °C until they reached the developmental stage required for each assay. For aging studies, day 1 adults were placed on plates containing 50 μM FUDR (RPI) and moved to fresh FUDR plates every other day until evaluation.

### Short-term associative learning

Short-term associative learning was performed as previously described (9) with minor modifications. Animals were age-synchronized via bleaching and allowed to hatch overnight in S-basal supplemented with 0.05% PEG-8000. Approximately 250 L1 larvae per condition and replicate were transferred onto 6-cm NGM plates and incubated at 20 °C until they reached the appropriate developmental stage for the assay. For aging studies, day 1 adults were placed on plates containing 50 μM FUDR (RPI) and moved to fresh FUDR plates every other day until evaluation. The day before the experiment, animals were transferred to plates dried for 2 h in a laminar flow hood. On the day of the experiment, conditioned animals were exposed to butanone by streaking 5 μL of a 10% (v/v) butanone solution in ethanol on the underside of the lid of a 6-cm plate, and incubated in a humidified chamber for 60 min at 20 °C. After conditioning, animals were washed off the plate using 1 mL of S-basal + 0.05% PEG-8000 and transferred into 1.5-mL tubes to settle for approximately 2 minutes. A 20 μL aliquot of the settled pellet was then placed at the origin of a 10-cm chemotaxis plate containing butanone and ethanol. Chemotaxis plates were prepared by spotting 2 µL of 2% sodium azide onto the locations designated for butanone or ethanol. Plates were left for approximately 10 minutes to allow the sodium azide to absorb into the agar. Subsequently, 2 µL of ethanol or 10% butanone (v/v) were added to the respective spots on each plate. The same procedure was followed for non-conditioned control animals. Excess moisture at the deposition site was gently absorbed using a folded tissue to release animals onto the plate for chemotaxis. After 1 h of chemotaxis at 20 °C, plates were transferred to a 4 °C cold room to immobilize the animals until scoring.

The learning index was calculated as the difference between the chemotaxis index (for butanone-conditioned animals) and the naive index (for non-conditioned animals), where each index corresponds to [(number of animals in butanone − number of animals in ethanol) / total number of animals not remaining at the origin].

### Thrashing

Thrashing was assessed by counting the number of body bends made by each animal in 20 µL of M9 buffer over a 30-second period. The assay was performed on at least 20 animals per condition. Animals were age-synchronized via standard hypochlorite treatment of gravid hermaphrodites to isolate eggs, which were allowed to hatch overnight in S-basal supplemented with 0.05% PEG-8000. Approximately 150 synchronized L1 larvae per condition were transferred to 6-cm NGM plates and incubated at 20 °C until they reached the developmental stage required for each assay. For aging studies, day 1 adults were either transferred to plates containing 50 μM FUDR (RPI) and moved to fresh FUDR plates every other day until evaluation or transferred daily to fresh plates without FUDR using a platinum wire.

### Movement

Spontaneous movement was assessed by counting the number of body movements (sinusoidal oscillations and spontaneous reversals) over 30-second intervals by direct observation under a stereomicroscope. Animals were prepared for evaluation as in the thrashing assay.

### Osmotic stress resistance assay

For the osmotic stress resistance assay, 30 synchronized animals at day 1, 5, or 10 of adulthood were manually transferred using a platinum wire to NGM plates containing high salt concentration (400 mM NaCl) and seeded with *E. coli* OP50. Plates were incubated at 20 °C, and survival was assessed after 24 hours by gently prodding with a platinum wire. Animals that failed to respond were scored as dead. Treatment with DMSO (vehicle) or 10 nM ADIOL was initiated at the L4 stage and discontinued at the time of transfer to the osmotic stress plates. To assess animals on days 5 and 10 of adulthood, they were transferred daily to fresh FUDR-free plates using a platinum wire until reaching the corresponding stages.

### Benzaldehyde chemotaxis

The benzaldehyde chemotaxis assay was performed as previously described (9). The benzaldehyde chemotaxis index was calculated as (number of animals in benzaldehyde - number of animals in ethanol)/(total number of animals). To prepare the animals for the assay, gravid hermaphrodites were age-synchronized via standard hypochlorite treatment to isolate eggs, which were allowed to hatch overnight in S-basal supplemented with 0.05% PEG-8000. Approximately 150 synchronized L1 larvae per condition were transferred to 6-cm NGM plates and incubated at 20 °C until they reached the developmental stage required for each assay. For aging studies, day 1 adults were placed on plates containing 50 μM FUDR (RPI) and moved to fresh FUDR plates every other day until evaluation.

### Lifespan

For lifespan assessment, animals synchronized by egg-laying instead of bleaching were grown until day 1 of adulthood on 6-cm NGM plates seeded with *E. coli* OP50. From that day onward, survival was monitored daily until no live adults remained. Animals were scored as dead if they failed to respond when gently prodded with a platinum wire. In all assays, animals were transferred to plates containing 50 μM FUDR from day 1 to day 8 of adulthood, being moved to fresh plates every other day. The only exception was the CR experiment, where *ad libitum* (AL) fed animals were transferred every other day to fresh plates seeded with *E. coli* OP50 using a platinum wire, while CR animals were transferred every other day starting from day 3 of adulthood to NGM plates without OP50, relying solely on the small amount of bacteria carried over with the platinum wire during transfer.

### RT-qPCR

Approximately 300 synchronized L1 larvae were transferred onto 6-cm NGM plates per replicate and allowed to develop to the L4 stage. Worms treated with DMSO (vehicle) or F17 were exposed to the compounds overnight prior to reaching the L4 stage. Total RNA was extracted from these animals using TRIzol reagent and purified with the Direct-zol RNA Miniprep Kit (Zymo Research), including on-column DNase treatment. First-strand cDNA was synthesized using the ProtoScript II First Strand cDNA Synthesis Kit (New England Biolabs) with random primers.

Quantitative PCR was performed using the SsoAdvanced Universal SYBR Green Supermix (Bio-Rad) on a C100 Touch™ Thermal Cycler CFX384™ Real-Time System (Bio-Rad), in 384-well plates. The reactions (10 µL final volume) contained 5 µL of SYBR Green Supermix, 0.5 µL of gene-specific primers (5 µM), 1 µL of cDNA (diluted 1:4), and 3.5 µL of nuclease-free water.

Cycling conditions followed a two-step protocol: initial denaturation at 95 °C for 3 min, followed by 40 cycles of 95 °C for 10 s and 60 °C for 30 s, ending with a melting curve analysis. Each sample was analyzed in technical triplicates, and appropriate negative controls were included. Threshold cycle (Ct) values were obtained using CFX Manager v3.1 software (Bio-Rad). Gene expression was normalized to the reference gene *tba-1*. Relative expression levels were calculated using the 2^−ΔΔCt^ method. Primer sequences are provided in **Supplementary Table 2**.

### Data visualization and statistics

Statistical analyses and graphical representations were performed using StataSE v12 (StataCorp LP, College Station, TX, USA) and GraphPad Prism 8.0.2 (GraphPad Software, San Diego, CA, USA). Normality was assessed using the Shapiro-Wilk test, except for feeding and thrashing data, for which the D’Agostino test was used due to the high number of repeated values. To enhance clarity, the choice between reporting the mean +/− standard deviation (SD) or the median +/− interquartile range (IQR), as well as the specific statistical test used, is indicated in each figure legend. The significance level (α) was set at 0.05, except for the log-rank test, where it was adjusted to 0.01 to reduce the risk of type I errors (false positives).

## DECLARATIONS

### Author contributions

Conceptualization, K.A., G.A.L., and A.G.-H.; methodology, A.G.-H., S.Y., and G.A.L.; formal analysis, A.G.-H.; investigation, A.G.-H., S.Y., S.K., A.Q.L., R.R.P.; resources, K.A.; data curation, A.G.-H.; validation, S.Y., S.K., and A.G.-H.; writing – original draft preparation A.G.-H., K.A.; writing – review & editing, K.A., A.G.-H., G.A.L., S.Y., S.K.; visualization, A.G.-H.; supervision, K.A.; funding acquisition, K.A.

## Acknowledgements

Some strains were provided by the Caenorhabditis Genetics Center (CGC), which is funded by NIH Office of Research Infrastructure Programs (P40 OD010440). This work was supported by NIH grants R01AG046400 and RF1AG068194 to K.A. Additionally, support was provided by the UCSF Program for Breakthrough Biomedical Research, funded in part by the Sandler Foundation.

