## Supplementary Information for "Dietary restriction promotes neuronal resilience via ADIOL"

**Supplementary Figure 1**

**Extended Figures 1 and 2**

**Supplementary Tables 1 and 2**

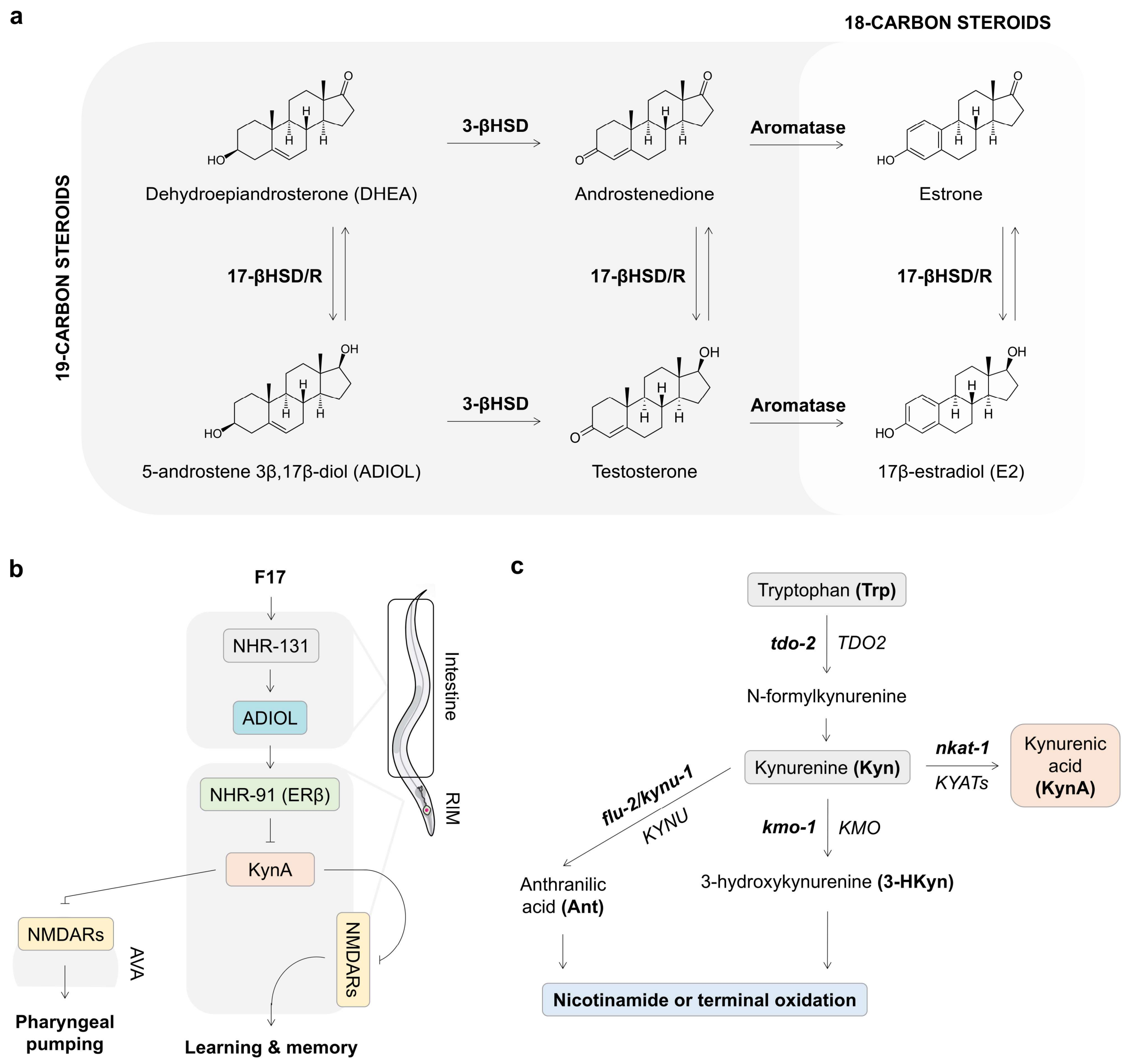


**Supplementary Fig. 1**. **a,** 5-androstene-3β,17β-diol (ADIOL) is a 19-carbon steroid produced from dehydroepiandrosterone (DHEA), which acts as an intermediate in the synthesis of other 19-carbon steroids such as testosterone, as well as 18-carbon steroids like 17β-estradiol (E2). **b,** F17 promotes learning and memory and pharyngeal pumping through KynA modulation. The compound F17 activates NHR-131 in the intestine, leading to an increase in ADIOL production. NHR-91 (ERβ) in the RIM neuron is required for the learning and pumping effects of ADIOL. ADIOL-induced decrease in KynA activates RIM and AVA neurons in NMDAR-dependent manner, which promotes learning and memory and pharyngeal pumping, respectively. **c,** Tryptophan metabolism pathway leading to kynurenic acid (KynA) production. Tryptophan is converted to kynurenine (Kyn) by the enzyme TDO2 (TDO-2). Kyn is then metabolized either into KynA via KYAT enzymes (NKAT-1) or further processed into nicotinamide or through terminal oxidation.

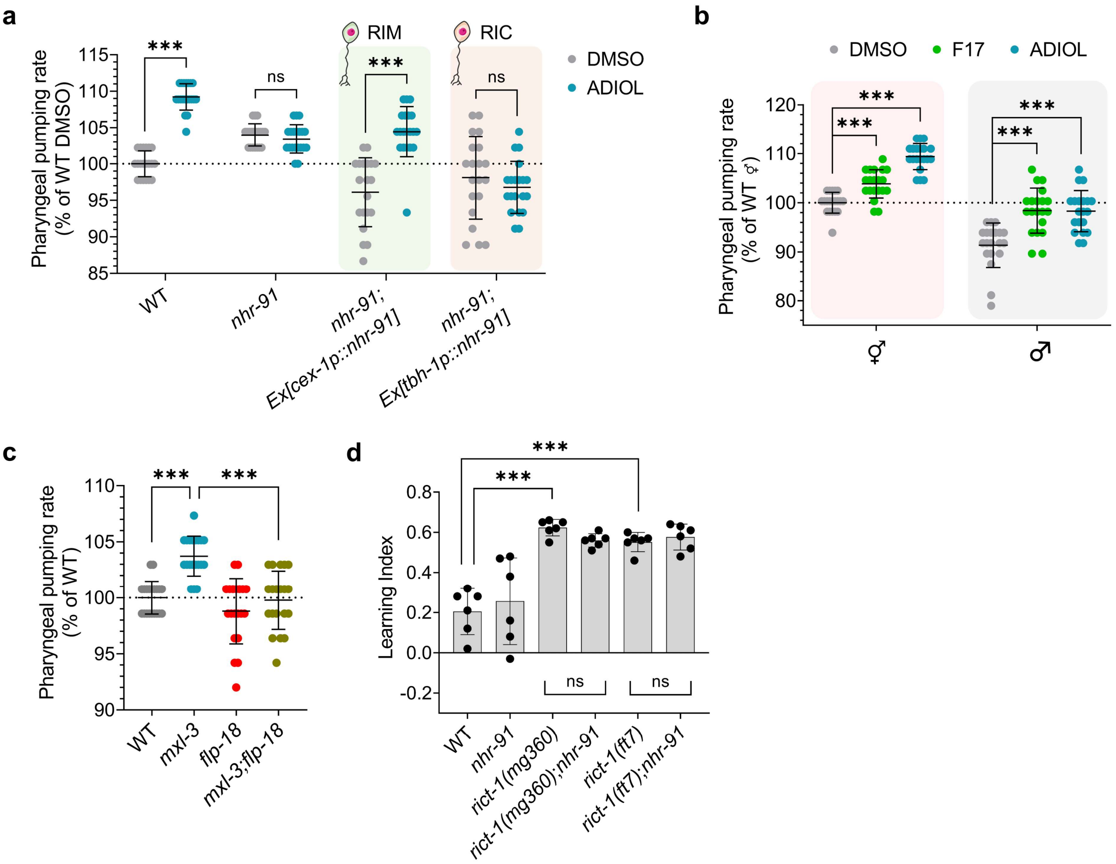


**Extended Fig. 1**. **a-c**, Pharyngeal pumping rate in day 1 adults, normalized to WT controls (n = 20 animals/condition; mean ± SD). (a) WT, *nhr-91*, and rescue strains (*nhr-91*;Ex[*cex-1p::nhr-91*,*unc-122::GFP*], and *nhr-91*;Ex[*tbh-1p::nhr-91cDNA::sl2::GFP*]) treated with DMSO or 10 nM ADIOL from the L4 stage. Statistics: t-test (***p < 0.001, ns: non-significant). (b) WT hermaphrodites and males treated with DMSO, 2.5 µM F17 or 10 nM ADIOL from the L4 stage. Statistics: one-way ANOVA with Bonferroni's correction (***p < 0.001). (c) WT, *mxl-3*, *flp-18*, and *mxl-3*;*flp-18* strains. Statistics: one-way ANOVA with Bonferroni's correction (***p < 0.001). **d**, Learning index of the CR models *rict-1(mg360)* and *rict-1(ft7)* lacking NHR-91 on day 1 of adulthood (n = 6; 25-194 animals/replicate). Mean ± SD shown. Statistics: one-way ANOVA with Bonferroni's correction (***p < 0.001, ns: non-significant).

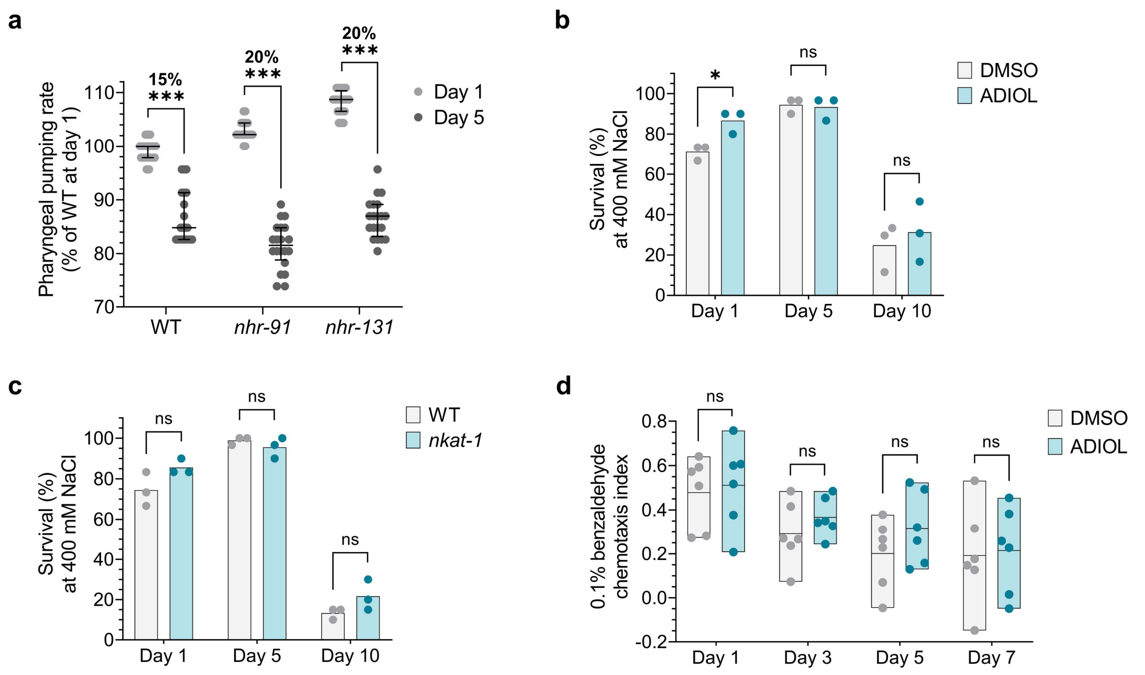


**Extended Fig. 2. a**, Pharyngeal pumping rate in WT, *nhr-91*, and *nhr-131* animals on days 1 and 5 of adulthood (n = 20 animals/condition). Median ± IQR values shown relative to day 1 WT adults. Animals were treated with 50 μM FUDR starting on day 1. Statistics: t-test (*p < 0.001). **b**,**c**, Survival after 24-hour exposure to 400 mM NaCl, normalized to WT controls on day 1. Mean ± SD shown (n = 3; ~30 animals/replicate). No FUDR was used in this assay. Statistics: t-test (***p < 0.001, **p < 0.01, ns: non-significant). (b) WT animals treated with DMSO or 10 nM ADIOL from the L4 stage. (c) WT and *nkat-1* animals. **d**, Chemotaxis index toward 0.1% benzaldehyde in WT animals treated with DMSO or 10 nM ADIOL on days 1, 3, 5, and 7 of adulthood. Treatment began at the L4 stage, with 50 μM FUDR from day 1. Mean ± SD shown (n = 3; ~38-95 animals/replicate). Statistics: t-test (ns: non-significant).

**Supplementary Table 1.** Strains used in this study.

| Strain name (in text) | Genotype |
|---|---|
| Wild-type/WT | WT |
| *nhr-91* | *nhr-91*(tm4713) X |
| *nhr-131* | *nhr-131*(tm1376) V |
| *nhr-91;*Ex[*cex-1p::nhr-91*] | *nhr-91*(tm4713) X; Ex[*cex-1p::nhr-91, unc-122::GFP*] |
| *nhr-91;*Ex[*tbh-1p::nhr-91*] | *nhr-91*(tm4713) X; Ex[*tbh-1p::nhr-91cDNA::sl2::GFP*] |
| *daf-2* | *daf-2*(e1370) III |
| *daf-2;nhr-91* | *daf-2*(e1370) III; *nhr-91*(tm4713) X |
| *daf-2;nhr-131* | *daf-2*(e1370) III; *nhr-131*(tm1376) V |
| *mxl-3* | *mxl-3*(ok1947) X |
| *mxl-3;nhr-91* | *mxl-3*(ok1947) X; *nhr-91*(tm4713) X |
| *mxl-3;nhr-131* | *mxl-3*(ok1947) X; *nhr-131*(tm1376) V |
| *flp-18* | *flp-18*(gk3063) X |
| *mxl-3;flp-18* | *mxl-3*(ok1947) X; *flp-18*(gk3063) X |
| *rict-1(mg360)* | *rict-1*(mg360) II |
| *rict-1(mg360);nhr-91* | *rict-1*(mg360) II; *nhr-91*(tm4713) X |
| *rict-1(ft7)* | *rict-1*(ft7) |
| *rict-1(ft7);nhr-91* | *rict-1*(ft7); *nhr-91*(tm4713) X |
| *cyp-44A1* | *cyp-44A1*(ok216) II |
| *cyp-13A4* | *cyp-13A4*(tm7443) II |
| *F12E12.11* | *F12E12.11*(ft1005) II |
| *nkat-1* | *nkat-1*(ok566) X |
| *kmo-1* | *kmo-1*(tm4529) V |
| *eat-2* | *eat-2*(ad465) II |
| *eat-2;nhr-91* | *eat-2*(ad465) II; *nhr-91*(tm4713) X |

**Supplementary Table 2.** qPCR primers used in this study.

| Gene | F/R | Sequence | Tm | Genomic length | Mature length |
|---|---|---|---|---|---|
| ***cyp-44A1*** | F | CTCGAATCTGCTGGTCAAATA | 60 | 200 | 156 |
| | R | TGTTGAGAACAGACGGAATATC | 60 | | |
| ***cyp-13A4*** | F | GCTCTACGAATGTACCCTTTAG | 60 | 214 | 120 |
| | R | AAGTGTCCATGTATCCACTTG | 60 | | |
| ***cyp-33C2*** | F | CCCGTTCTCAGTTGGAAA | 59 | 197 | 146 |
| | R | GGGCTCCATTGCTCTTATC | 60 | | |
| ***cyp-33C5*** | F | GAATGAGACACTTGGTGGAG | 60 | 256 | 207 |
| | R | GGCTTTCCTCGGAATCAAA | 60 | | |
| ***cytb-5.1*** | F | CTGACGCTAGGCATATGAAG | 60 | 166 | 112 |
| | R | TTATCCTGTTCGGTGGTAGA | 60 | | |
| ***F12E12.11*** | F | AGTCTACACAGGATTCGGAG | 60.4 | 400 | 127 |
| | R | GGCAATTTCTATGGGCTGAG | 60.8 | | |
| ***F25D1.5*** | F | GCTGTGATCGAAATGACTCAG | 60.9 | 203 | 156 |
| | R | GCATCTGGTGTATTGGTCAAG | 61 | | |
| ***R05D8.9*** | F | CGGGTTTGGAGAAGCTATG | 60 | 184 | 141 |
| | R | TCAGCCAAGAACGCAATAA | 60 | | |
| ***tba-1**** | F | ACACTCCACTGATCTCTGC | 60.9 | 174 | 129 |
| | R | CAGCCATGTACTTTCCGTG | 60.5 | | |

*Reference gene
